# LactoSpanks: a collection of IPTG inducible promoters for the commensal lactic acid bacteria *Lactobacillus gasseri*

**DOI:** 10.1101/2023.07.13.548755

**Authors:** Elsa Fristot, Guillaume Cambray, Jerome Bonnet

## Abstract

Lactic acid bacteria (LAB) are important for many biotechnological applications, such as bioproduction and engineered probiotics for therapy. Inducible promoters are key gene expression control elements, yet those available in LAB are mainly based on bacteriocin systems and have many drawbacks, including large gene clusters, costly inducer peptides and little portability to *in vivo* settings. Using *Lactobacillus gasseri*, a model commensal bacteria from the human gut, we report the engineering of Lactospanks promoters (Pls), a collection of variable strength inducible promoters controlled by the *LacI* repressor from *B. subtilis* and induced by isopropyl β-D-1-thiogalactopyranoside (IPTG). We first show that the Phyper-spank promoter from *Bacillus subtilis* is functional in *L. gasseri*, albeit with substantial leakage. We then construct and screen a semi-rational library of Phyper-spank variants to select a set of four IPTG-inducible promoters that span a range of expression levels and exhibit reduced leakages and operational dynamic ranges (from ca. 9 to 28 fold-change). With their low genetic footprint and simplicity of use, Lactospanks will support many applications in *L. gasseri*, and potentially other lactic acid and gram-positive bacteria.

## MAIN TEXT

Lactic acid bacteria (LAB) encompass a group of disparate microorganisms present in various ecological niches from fermented food to mammalian microbiota^1^. LAB are gram-positive, rod-shaped, non-spore forming, acido-tolerant and facultative anaerobic bacteria that share the ability to convert carbohydrates to lactic acid^2^. Due to their GRAS (generally recognized as safe) qualification, they are of great interest for biotechnological applications, including fermentation^3^, production of food preservatives^4^ and, more recently, engineered probiotics for therapy^5^. Many of these applications primarily rely on the precise control of gene expression.

Several systems have been deployed to control gene expression in LAB. While initial approaches leveraged constitutive promoters from housekeeping genes (*e*.*g*. PslpA and Ppgm from *L. acidophilus*)^6^, inducible gene expression systems provide greater versatility for rapid prototyping, controllable production of physiologically costly proteins or sensitive *in vivo* applications. The main induction systems used in lactic acid bacteria are based on bacteriocin-^7,8^, stress-^9^ or carbohydrates-inducible systems^10^. The Nisin inducible pNICE system is the reference, but it involves a large gene cluster that limits plasmid real estate and reduces transformability. Furthermore, the system can impose a burden on the host cell, as it requires expression of a membrane receptor that can alter bacterial physiology and the induction with Nisin peptide, a bacteriocin which can be toxic in some expression strains^11^.

In widely used and studied model organisms, such as *Escherichia coli*, a wide variety of inducible systems with low genetic burden have been developed. Examples include the Ptet system induced by anhydrotetracycline, the Pbad system controlled by arabinose, or the Plac system inducible with lactose or its non metabolisable analog isopropyl β-D-1-thiogalactopyranoside (IPTG)^12^. These inducible systems are extensively applied to control the levels and timing of gene expression in liquid culture and *in vivo* applications. Contrasting with those currently used in LAB, these classical systems rely on the expression of a single repressor gene, which involves lower genetic footprint, complexity and burden, and consequently improved portability. They also respond to cheap inducers that are non toxic and, for some of them, non-metabolizable (*e*.*g*. aTc or IPTG). Although some Plac systems have been reported in lactic acid bacteria, their use has remained anecdotal due to limited dynamic ranges (at most four-fold) and induction restricted to lactose instead of non-metabolizable analogs^13–15^. Optimization of these systems to reduce leakage while maintaining strong expression upon induction is thus essential to improve their utility. Such systems (based on *Tet, LacI*, or *T7*) have been successfully developed in model gram-positive bacteria, such as *B. subtilis*, and several have proven functional in various lactobacilli (*e*.*g*. PSfrA)^16^.

We are interested in developing inducible systems for *Lactobacillus gasseri ATCC33233*^17^, an easily manipulated strain that is naturally present in the vaginal and colorectal microbiota and has the potential to be engineered as a boosted probiotic, with existing proof of principles for efficient mucosal vaccination^18^. To develop reliable inducible promoters for the precise engineering of protein delivery by this bacteria, we chose to use the Phyper-spank promoter, an inducible system developed for *B. subtilis* IPTG induction derived from the *spac* system carrying the bacteriophage promoter SP01 modified with lac operator and an *E. coli* lac repressor^19^.

As Phyper-spank was obtained from a *B. subtilis* shuttle vector, we first transferred the Phyper-spank to drive *sfGFP* expression along with its *LacI* repressor on the broad range lactobacilli shuttle vector pTRKH2 carrying the pAMB1 high-copy replication origin for gram-positive bacteria^20^, and transformed it into *L. gasseri* for characterization assays (**Figure 1A**). This construct showed first to be efficiently induced in the presence of IPTG but shows a sizable basal activity —about 5 times the background fluorescence of cells devoid of plasmid grown in the same conditions—thus indicating the *B. subtilis* promoter to be functional but substantially leaky in this strain (**Figure 1B and Table 1**). When induced with 100μM IPTG, Phyper-spank produced a strong response with a 16-fold increase in measured fluorescence levels.

**Table 1:** Phyper-spank and Pls variants characteristics measured after 8 hours of growth in the absence and presence of IPTG (100μM). Leakage is the median fluorescence intensity in the non-induced state minus median autofluorescence of negative control cells (without plasmid). The fold change (or dynamic range) is the ratio of median fluorescence intensities measured in induced over uninducedconditions. The swing isthe median fluorescence intensities measured in induced minus that measured in uninduced conditions. The EC50 is the half-maximal effective concentration derived from the fit shown in Figure 2C.

|  | Phyp | Pls1 | Pls2 | Pls3 | pLs4 |
| --- | --- | --- | --- | --- | --- |
| **% switched cells at 8 hours** | 86 % | 79 % | 71 % | 78 % | 75 % |
| **Fold change** | 16,15 | 8,98 | 20,82 | 28,10 | 18,89 |
| **Swing (A.U.)** | 71653,33 | 8649,33 | 25174 | 45502,33 | 52932,33 |
| **EC50 (IPTG M)** | $8,2 \pm 0,3 \times 10^{-6}$ | $29,4 \pm 1 \times 10^{-6}$ | $15,3 \pm 0,8 \times 10^{-6}$ | $17,2 \pm 0,8 \times 10^{-6}$ | $10 \pm 0,4 \times 10^{-6}$ |
| **Leakage (A.U.)** | 3933,33 | 287,33 | 473,66 | 882,66 | 2162,66 |

**Figure 1.**
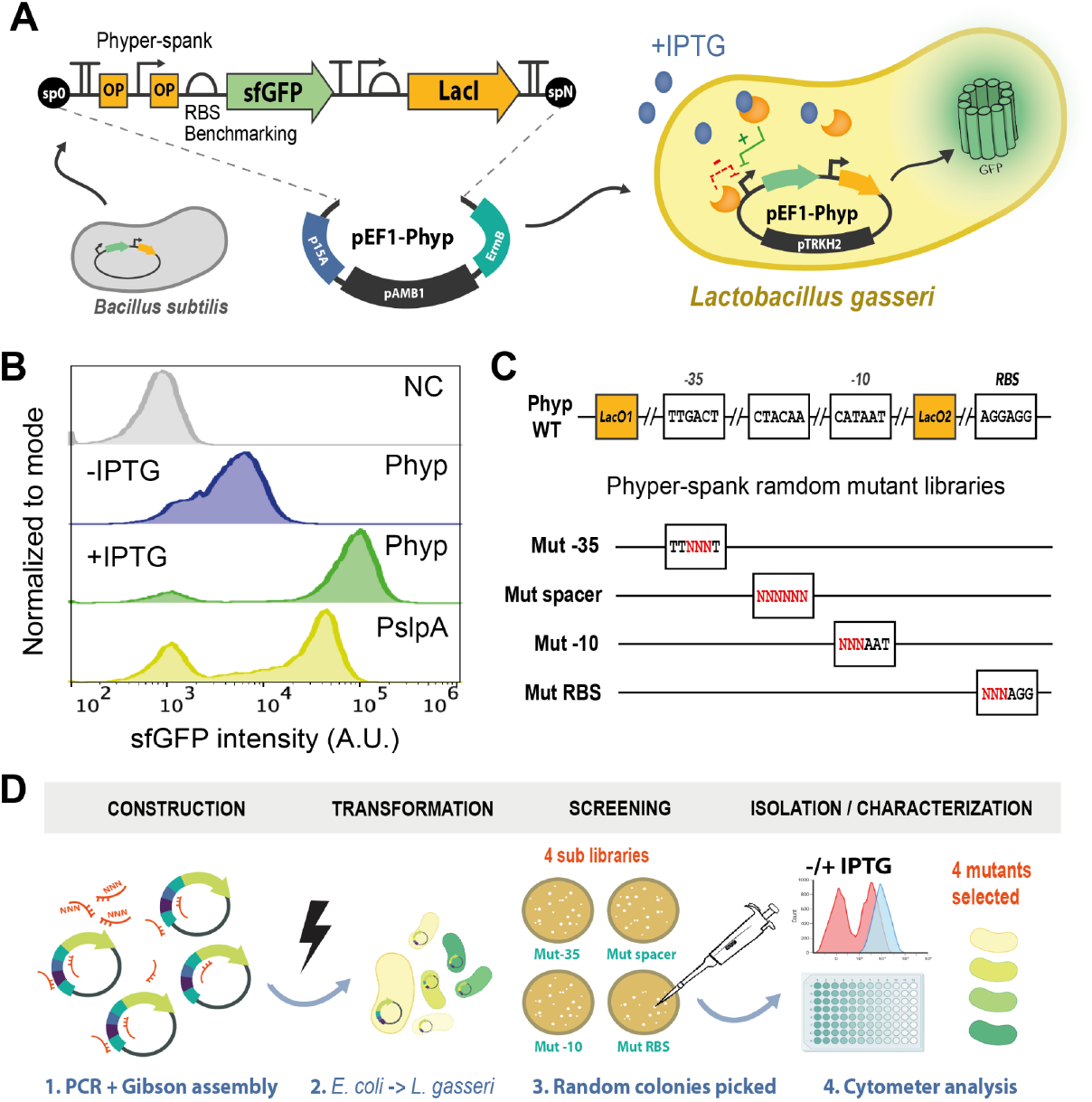
Generation and characterization of IPTG responsive promoter in *L. gasseri*. **A**. Transfer of Phyper-spank (Phyp) from *B. subtilis* to *L. gasseri*. Phyp was cloned in the shuttle vector pTRKH2 and transformed into *L. gasseri*. The plasmid map is shown in Supplementary Figure S1. **B**. Representative distribution of fluorescence intensities from Phyp operating in *L. gasseri* after 8 hours of growth in the absence (blue) or presence (green) of IPTG (100μM) as measured by flow cytometry. NC: negative control (grey), the WT *L. gasseri* without plasmid. PslpA (yellow): widely constitutive promoter used as a reference **C**. Schematic overview of four Phyper-spank regions independently targeted for random substitutions. Mutations are represented by red N characters in each sub-libraries,. **D**. Experimental workflow for generation, screening, and characterization of Phyper-spank variants: (1) Randomized promoters are generated by PCR using degenerated oligonucleotides, and inserted in pTRKH2 vectors by Gibson Assembly in *E. Coli*. (2) Plasmid clones are batch extracted and transformed into *L. gasseri*. (3) Colonies are randomly picked, grown in the absence or presence of IPTG (100μM) and analyzed by flow cytometry. (4) Four variants with improved dynamic ranges and graded expression levels are chosen for deeper characterization to form the Lactspank collection.

Considering the leakiness of the repressed Phyper-spank (referred as Phyp) and its otherwise strong intrinsic activity, we reasoned that improved dynamic ranges could be obtained by reducing the strength of the promoter. To achieve this, we separately introduced random nucleotide substitutions in three distinct regions known to modulate the activity of σ^70^ promoters to various extent : the -35 motif, the -10 motif, and the spacer between those two regions. To further investigate the potential impact of translation on the response, we also generated a fourth library targeting the ribosome binding site (RBS). **(Figure 1C)**. Each sub-library was generated using degenerate oligonucleotides, cloned in *E. coli*, transformed in *L. gasseri* and screened by randomly picking 12-40 clones and coarsely evaluating their response to induction in a single replicate **(Figure 1D)**.

All isolated clones exhibited reduced leakage due to decreased gene expression caused by the mutated promoter regions (**Supplementary Figure S3**). Broadly speaking, mutations in the -10 motif or RBS resulted in strong reductions in reporter expression, while mutation in the -35 and between the -35 and the -10 regions leads to a more moderate decrease. Based on improved dynamic range (ratio between the fluorescence intensities of the fully activated state vs the non-induced state).. Based on these initial results, we chose 22 promising clones (10, 9, 1 and 2 for the -35, spacer, -10 and RBS regions, respectively) to perform three additional replicates induction experiments (**Supplementary Figure S4)**.

We categorized these 22 clones into four levels of induced expression, then chose the clone with maximal dynamic range from each bin, which yielded four final variants Pls1 to 4. We compared their expression profiles with the original Phyper-spank construct under saturating induction with 100 μM IPTG (**Figure 2A and 2B**). Depending on the mutation, these variants showed a 2 to 10 fold reduction in basal leakage while the maximal induced expression is comparatively less affected. For example, Pls4’s leakage is reduced by 50%, but its induced expression is only reduced by 25%. Across all variants, the reduction in leakage tends to be twice as marked as the reduction in induced expression, which results in better dynamic ranges, with a fold change up to twice as high as that of Phyper-spank.

**Figure 2:**
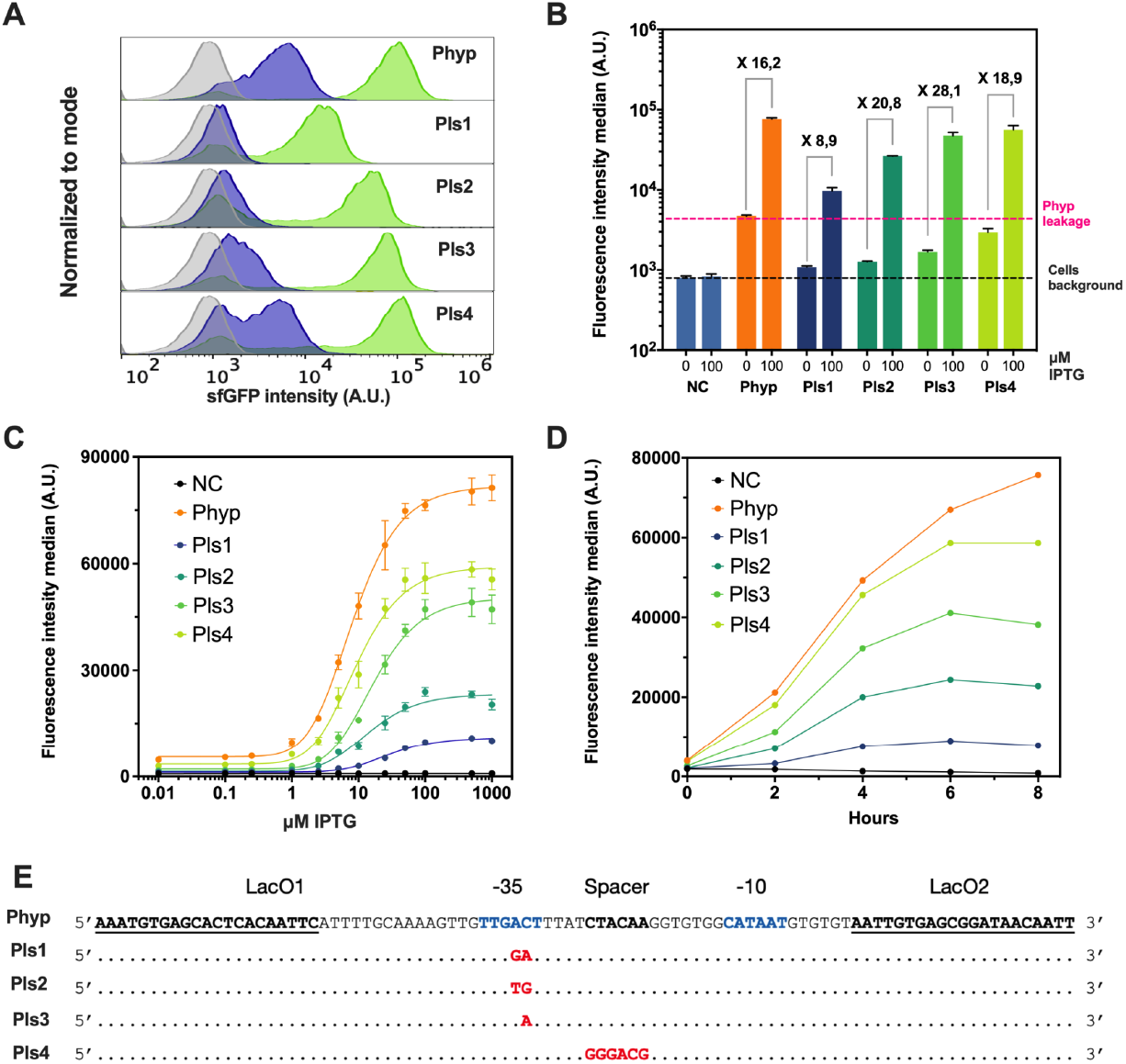
Characterization of the Lactospank promoter collection. **A**. Representative distribution offluorescence intensities from Phyper-spank and the four Pls variants after 8 hours of growth in the absence or presence of IPTG (100μM), as measured by flow cytometry. **B**. Mean of the median *sfGFP* fluorescence intensities for the different variants after 8 hours of growth in the absence or presence ofIPTG (100μM). Means are calculated over three replicate cultures grown on different days. Error bars: +\-standard deviation. Resulting dynamic range for each variant is shown above the corresp[onding bars.. NC: negative control (grey), the WT *L. gasseri* without plasmid. **C**. Response of the different variants to increasing concentrations of IPTG after 8 hours of growth.. Data points show the means of the median fluorescence intensities for three replicates cultures grown on different days. Error bars: +\-SD. NC: negative control, the WT *L. gasseri* without plasmid. Data is color-coded for the different strains, as shown. **D**. Kinetic response of the different variants after induction by IPTG (100μM). Shown are the medians of *sfGFP* fluorescence intensities measured by flow cytometry. NC: negative control, the WT *L. gasseri* without plasmid. Data is color-coded for the different strains, as shown. **E**. Sequences of the four Pls variants and comparison with Phyper-spank original sequence. Mutations are shown in red.

We next proceeded to measure the variants’ response to increasing concentration of IPTG, ranging from 10nM to 1mM (**Figure 2C, supplementary figure S5**). The four variants cover a wide range of maximum expression levels, ranging from 25 to 75% that of Phyper-spank (**Figure 2C and table 1**). The half maximal effective concentration (EC50) of IPTG derived from these titration curves tends to increase with reduced promoter strength (**Table 1**). We thus improved promoter fold changes compared to wt Phyper-spank, with Fold changes from 16 for the wild type version up to ∼28 for Pls3.

Promoters activation kinetics were then assessed (**Figure 2D**). The number of cells in the ON state increased with time, with the fraction of switched cells ranging from 21% for Pls1 up to 67% for Pls3 as early as two hours post induction, and a maximum reached after 6 post induction, with 73∼77% of switched cells for the Pls variants. Cells harboring Phyp show a switch from 62% 2 hours post-induction to 86% at 6 hours (**Table 1, supplementary table 1, supplementary figure S6**). A sizable fraction (from 14% for pHyp to 29% for Pls3) of the cell population maintains background fluorescence level, indicating the absence of reporter signal. We observed this behavior with all promoters we tested, including constitutive promoters of varying strength (*e*.*g*. reference promoter pSlpA on figure 1B).

The Pls1-4 variants were sequenced (**Figure 2D, and supplementary data**). The lowest expression mutants Pls1 and Pls2 showed distinct substitutions of two nucleotides at the same location of the -35 region while Pls3 showed a single mutation in the same region, supporting that the -35 motif is a strategic target to fine-tune gene expression. The strongest variant, Pls4 showed a 6 nucleotides substitution in the spacer region. Despite it harboring the largest number of mutations, the spacer variant had the smallest reduction in expression (**supplementary Figure S3D**).

During the course of our project, we identified limitations with the use of the *sfGFP* reporter in lactic acid bacteria. In the time-course experiment shown in figure 2D, the Phyper-spank construct did not reach saturation after 8h. In contrast, the fluorescence signal all of the weaker Pls variants saturated in 6h and showed a slight decrease at 8 hours post-induction, despite the stability of the *sfGFP* reporter protein. As with the fraction of non-expressing cells, the mechanism underlying this observation is unknown but may be linked to the acidification of the cytosol over time. In this perspective, the decrease in fluorescence may reflect the pH effect on the protonation of the *sfGFP* chromophore^21^, which would have a more prominent impact as the protein is expressed at lower levels. A similar observation was previously made in *L. casei*^15^.

The Lactospank collection provides a suite of four promoters (Pls1-4) with characteristics that can fit different user requirements. For instance, Pls3 demonstrates reduced leakage while reaching two-third the maximum expression level of the original Phyper-spank (**Table 1**), which should be useful for the majority of applications. Phyp and Pls2-4 Pls3 and Pls4 as well as the Phyp showed higher expression than the widely used strong constitutive promoter PslpA from *Lactobacillus acidophilus*, which placed them well for strong protein expression applications. In contrast, Pls1 provides lower expression levels, with no observable leakage, and will be suitable for expressing more toxic and complex compounds that require tight repression.

In Pls2-4, higher dynamic range was achieved through reducing basal leakage at the expanse of maximal induced expression with respect to the original Phyper-spank. Other approaches could be investigated to maintain or increase maximal expression, including tuning the expression of the LacI repressor, modulating the position and number of Lac operators, or using more advanced circuit topologies^22,23^.

A variety of orthogonal tools enabling fine tuning of gene expression are critically needed for lactic acid bacteria. Following this work, other low footprint inducible systems could be developed for these species. Some existing systems could be transposed from other species as we did. Alternatively, new inducible systems could be discovered in the wide realm of LAB and repurposed. Improved control will support the implementation of complex genetic circuits such as logic gates, oscillators, and memory systems, and allow synthetic biologists to unleash the potential of lactic acid bacteria as programmable agents for industrial, environmental, and medical applications.

## METHODS

### Plasmids

The pTRKH2 plasmid was akind gift from Rodolphe Barrangou & Todd Klaenhammer (Addgene plasmid #71312). pTRKH2 is a theta replicative plasmid carrying a pAMβ1 origin (high copy number), a p15A *E. coli* replication origin (low copy number) and an erythromycin resistance gene *Erm(B)*. The Phyper-spank-sfGFP sequence used in this study originated from our lab^24^. The Phyper-spank device and the pTRKH2 were PCR-amplified using the Q5 polymerase (NEB) and cloned by Gibson assembly^25^ in *E. coli* NEB 10 beta chemocompetent cells. The resulting plasmid pEF1-Phyperspank-sfGFP was sequenced and transformed in electrocompetent *L. gasseri* cells. The pSlpAs-sfGFP cassette was synthesized as a linear DNA fragment(Twist Bioscience) and cloned by Gibson assembly in pTRKH2. All constructs are flanked by standard spacers to facilitate downstream cloning via Gibson assembly. All plasmids are available from Addgene (pEF1-Phyper-spank-sfGFP: #205747, pEF1-Pls1-sfGFP: #205748, pEF1-Pls2-sfGFP: #205749, pEF1-Pls3-sfGFP: #205750, pEF1-Pls4-sfGFP: #205751). Primers are listed in Supplementary Table 2.

### Library generation

Phyper-spank variant libraries were constructed by amplifying the vector pEF1-Phyperspank-sfGFP with two set of primers to insert mutation in promoter region and reassemble the vector back using Gibson assembly in *E. coli* NEB 10 beta chemocompetent cells. Primer sequences are listed in Supplementary Table 2. Transformed cells were plated on LB agar plate with erythromycin (150μg/mL; E6376 Sigma-Aldrich) for selection. After overnight growth, all resulting colonies were scrapped from the plates and resuspended in 2mL LB with antibiotics. The four resulting cultures were grown overnight and subjected to plasmid extraction by miniprep. The plasmid libraries were then transformed into *L. gasseri* electrocompetent cells. After screening, plasmids clones were purified from *Lactobacillus gasseri* using the QIAprep spin Miniprep kit (Qiagen) with addition of 100μg/mL of Lysozyme and 5U/mL of Mutanolysin to buffer P1 with an incubation time of 30 minutes prior to lysis buffer addition, and sequence-verified by Sanger sequencing (Eurofins Genomics, EU).

### Strains and cell culture

*E. coli NEB10* beta was obtained from NEB. Cells are grown in LB media supplemented with Erythromycin (E6376 Sigma-Aldrich) at 150 μg/mL, as needed.*Lactobacillus gasseri* ATCC 33323 was obtained from LGC STANDARDS. ells are grown in liquid MRS medium (De Man, Rogosa and Sharpe, DIFCO) supplemented with erythromycin at 7 μg/mL, as needed. Growth is performed without agitation at 37°C in anaerobic conditions using CO2 incubator Eppendorf (10% CO2). MRS composition is as follow: peptone proteose: 10 g/L, beef extract: 10 g/L, yeast extract: 5.0 g/L, dextrose: 20 g/L, polysorbate 80: 1.0 g/L, ammonium citrate: 2.0 g/L, sodium acetate: 5.0 g/L, magnesium sulfate: 0.1 g/L, manganese sulfate: 0.05 g/L, dipotassium phosphate: 2.0 g/L. Agar plates are obtained by adding 1% agar to the appropriate liquid medium.

### Electrocompetent cells preparation

A 50 mL culture of *L. gasseri* cells was grown overnight. 98 mL of fresh MRS medium was then inoculated with 2 mL of the overnight culture and incubated to an OD of 0.5–0.6 (ca. 4–5 hours). Cells were harvested by centrifugation at 3,500 rpm for 5 min and the supernatant was discarded. From this step on, cells were always maintained on ice or at 4°C in refrigerated centrifuges. Pelleted cells were then resuspended in 15 mL ice-cold 3X SMEB electroporation buffer (1X Sucrose Magnesium Electroporation Buffer: 298 mM sucrose, 1 mM MgCl2 in cold sterile water), centrifuged at 5,000 rpm for 10 min and the supernatant was discarded. This procedure was repeated twice. Cell pellets were then concentrated in 1.0 mL of ice-cold 3X SMEB buffer and 200 μL aliquots were used directly for electroporation up to one hour after preparation or placed at -80°C for long term storage.

### Electroporation procedure

A GenePulser X Cell apparatus (Bio-rad Laboratories, Richmond, CA) was used for electroporation of *L. gasseri*. 200 μL competent cells were thawed on ice, transferred in a 0.2 cm electroporation cuvette (Bio-rad Laboratories, Richmond, CA) after addition and gentle mixing of 1 μg of DNA (up to 5uL), and incubated on ice for 5 minutes. After drying the electrodes with a clean paper wipe, cells were electroporated with a tension of 1.8 V, a resistance of 600 Ω and a capacitance of 25 μF. Successful electroporations usually showed a time constant of 8.0–11 ms. Directly after, cells were gently resuspended in 800 μL MRS medium pre-heated at 37°C and transferred to culture tubes for a 2 hours recovery at 37°C without shaking in anaerobic incubator. 100 μL of the resulting culture were then spread on MRS agar plate (1% agar) supplemented with 7μg/mL erythromycin using glass beads (0.4 cm diameter).

### Induction assays

Overnight pre-cultures were started from single colonies on agar plates. The next day, saturated cell cultures (OD600∼2) were diluted 1:100 in 400 μL MRS in 96 deepwell plates and incubated up to eight hours at 37° without agitation. IPTG was added at the dilution stop as needed. 10 uL from these culture were resuspended in 96 well plates in 200 μL focusing fluid for flow cytometry analyses.

### Cytometry analysis

Flow cytometry was performed on an Attune NxT flow cytometer (Thermo Fisher) equipped with an autosampler and Attune NxT Version 2.7 Software. Experiments on Attune NxT were performed in 96-well plates with the following setting: FSC: 200 V, SSC: 380 V, and green intensity BL1: 460 V (488 nm laser and 510/10 nm filter). All events were collected with a cutoff at 20,000 single cell events. Every experiment included a negative control consisting in cells without plasmid grown in the same conditions to generate the gates. See**Supplementary Figure S2** for the gating strategy. Data were analyzed using Flowjo (Treestar, Inc) and Prism (GraphPad).

### Data processing

Each experiment was replicated three times on three different days. For each strain, data are summarized as the mean and standard deviation of the median of the *sfGFP* fluorescence intensity across these three replicates. Fold change and swing of each variant were calculated from the mean of the fluorescence median. The goodness of fit and the EC50 for each data set were calculated by applying nonlinear regression using the agonist versus response variable slope function in GraphPad Prism.

## Authors contribution

EF, JB, and GC designed the study. EF performed the experiments. EF, JB, and GC analyzed the results and wrote the paper.

## Associated content

Supporting Information is available.

-Supplementary materials

-Raw data

-DNA sequences

## Acknowledgements

We thank members of our groups and of the CBS for fruitful discussions and feedback. We thank M. Jules and Olivier Delumeau (INRA MICALIS, Jouy-en-Josas) for raising our interest in using *B. subtilis* parts in lactic acid bacteria. This work was supported by grants from INSERM, the Bettencourt-Schueller Foundation, the Canceropole Grand Sud-Ouest (JB), and a CNRS-ATIP (GC). EF is a recipient of Ph.D fellowships from the French Ministry of Research and the French Ligue contre le Cancer. The CBS acknowledges support from the French Infrastructure for Integrated Structural Biology (FRISBI) ANR-10-INSB-05-01.

