## Supplementary materials for "LactoSpanks: a collection of IPTG inducible promoters for the commensal lactic acid bacteria *Lactobacillus gasseri*"

pEF1-pHyperspank-sfGFP (8940 bp)

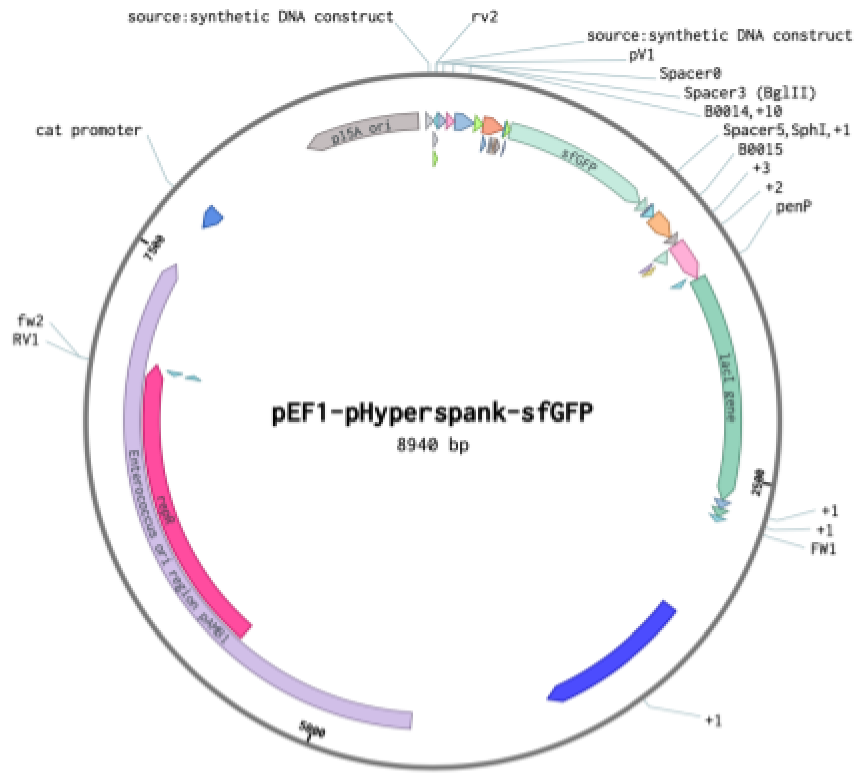


**Figure S1.** pEF1-Phyperspank-sfGFP plasmid map

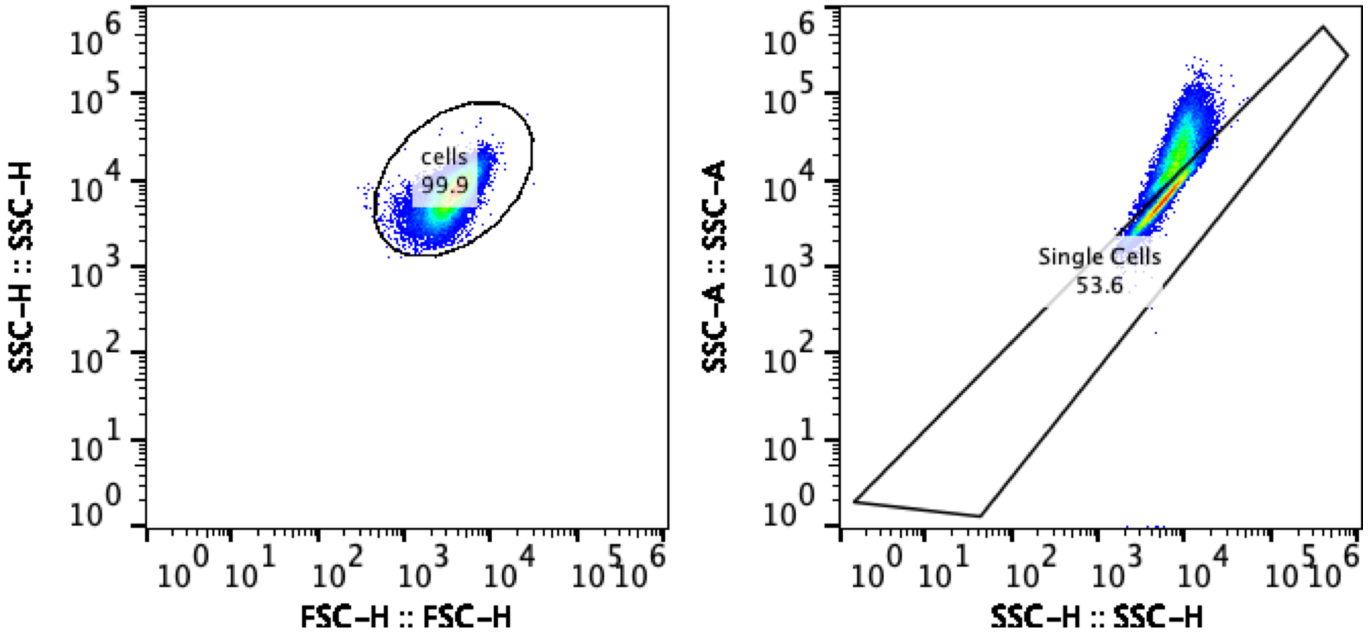


**Figure S2.** Figure exemplifying the gating strategy. Gates were designed based on FSC-H vs SSC-H scatter plots to remove debris from the analysis, and SSC-A vs SSC-H to doublet discrimination, left and right panel, respectively. The median of the sfGFP fluorescence was calculated from selected cells in the Single Cells gate.

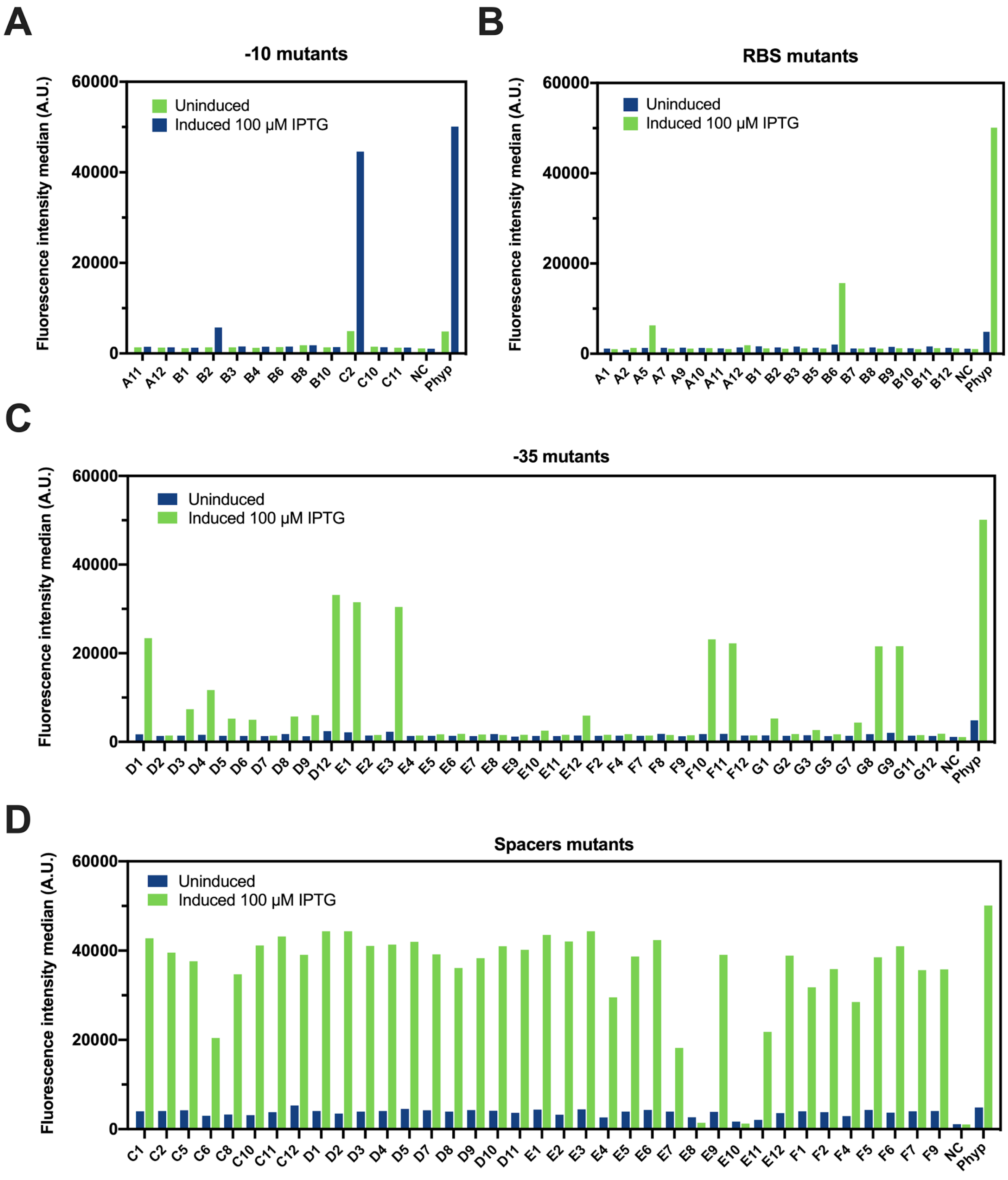


**Figure S3.** Screening of Phyper-spank mutants from the four sub-libraries by flow cytometry in *L. gasseri*. **A.** sfGFP fluorescence median of -10 mutant single clones. **B.** sfGFP fluorescence median of RBS mutant single clones. **C.** sfGFP fluorescence median of -35 mutant single clones. **D.** sfGFP fluorescence median of spacer mutant single clones. Induction with or without IPTG at 100 µM at hour 4 post induction.

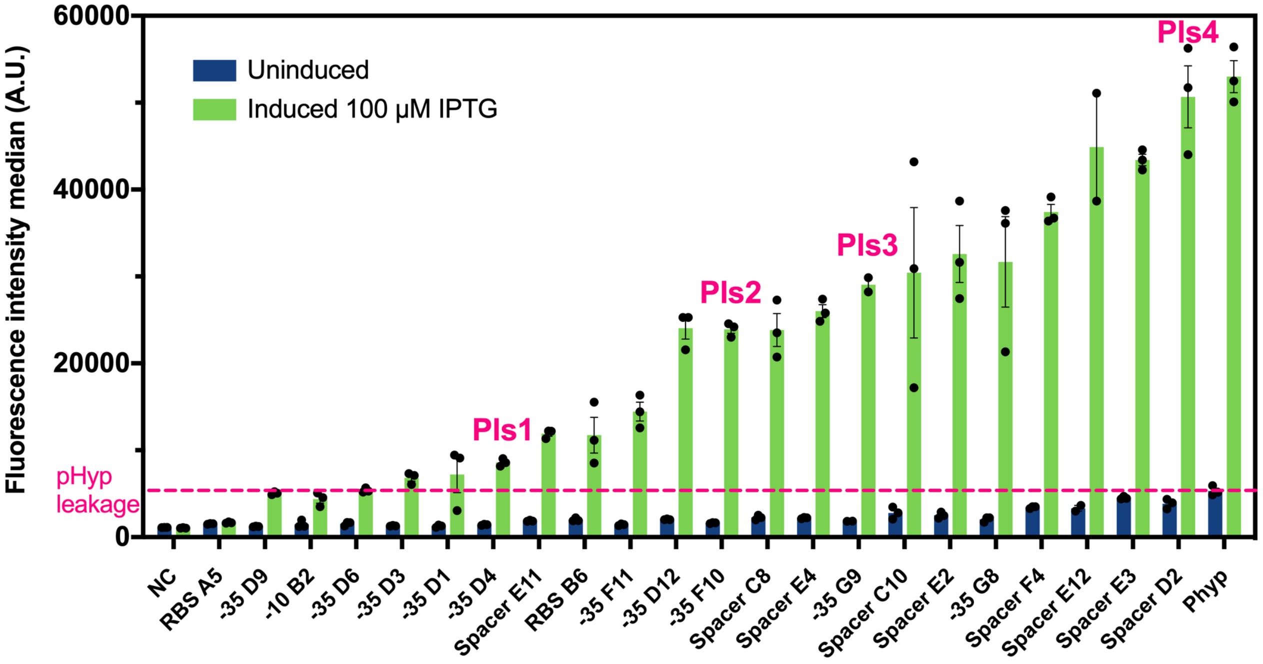


**Figure S4.** sfGFP fluorescence median measured by flow cytometry with or without 100µM IPTG at hour 4 post induction on the 22 mutants selected for their best dynamic range. Error bar represent the mean of three replicates with/without SEM .

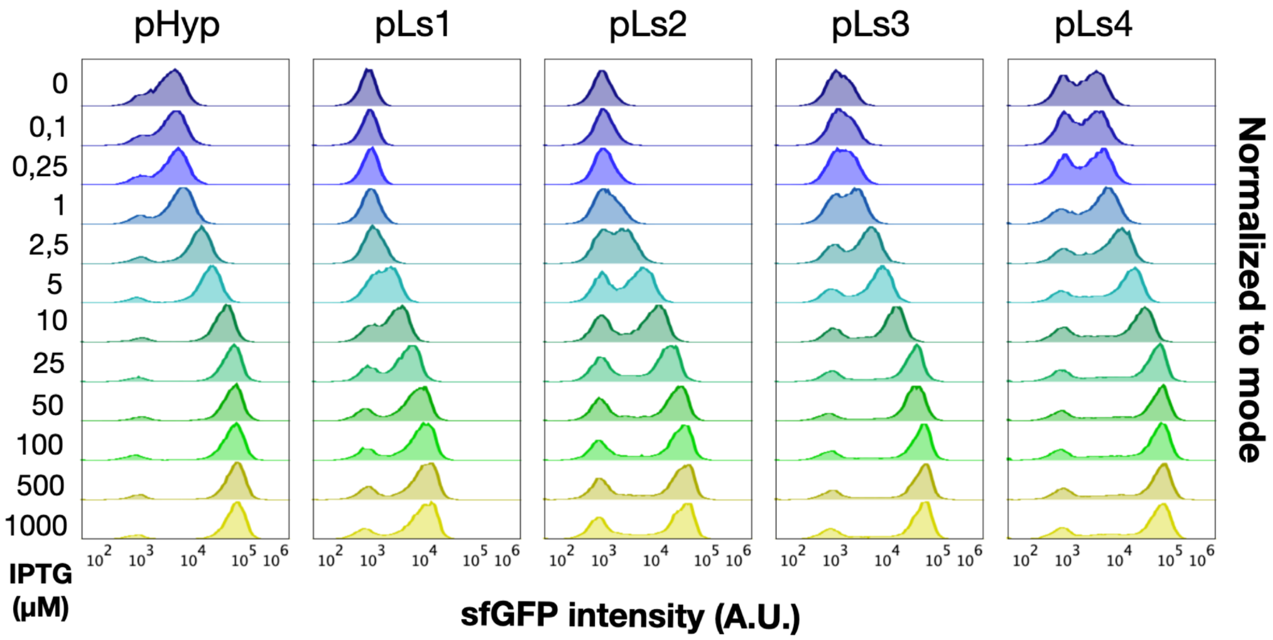


**Figure S5.** Representative flow cytometry histograms of the sfGFP fluorescence from Phyper-spank and the four pLs variants at hour 8 post induction with different level of IPTG.

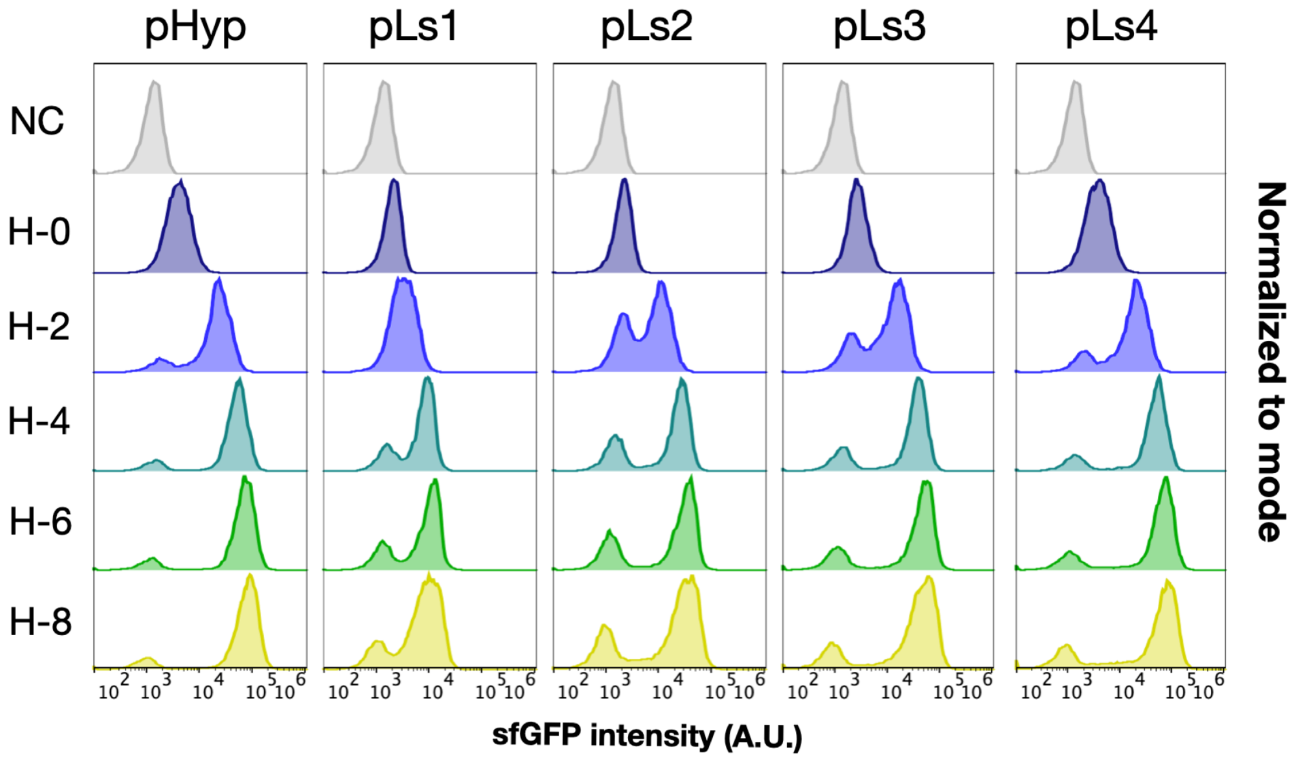


**Figure S6.** Representative flow cytometry histograms of the sfGFP fluorescence from Phyper-spank and the four pLs variants at four different time points after induction with 100μM of IPTG.

| | % shifted cells | | | | |
|---|---|---|---|---|---|
| | pHyp | pLs1 | pLs2 | pLs3 | pLs4 |
| H2 | 62,6 | 21,5 | 55 | 67,3 | 49,1 |
| H4 | 83,6 | 72,1 | 68,8 | 74,9 | 78 |
| H6 | 86,2 | 73,2 | 69 | 75,7 | 77,4 |
| H8 | 86,3 | 79,4 | 71,9 | 78 | 75,4 |

**Table S1. Pourcentage of switched cells at different time points after induction**

**Table S2. Oligonucleotides sequences**

| | Oligonucleotides | Sequence |
|---|---|---|
| pTRKH2 backbone amplification to insert pHyp | **pRpTRKH2-sp0** | aacagagtaagggtatccgaggctgcattaatgaatcggcca |
| | **pFpTRKH2-spN** | tagaatcgtgcttcagtaagagttgctgaacttttaaaacaagca |
| Amplification of pHyperspank device to insert in pTRKH2 | **Sp0_f** | CTCGGATACCCTTACTCTGTTGAAAAC |
| | **SpN_r** | TCTTACTGAAGCACGATTCTACTCGG |
| Insertion of mutation in RBS | **pFnnnRBS** | gctagcgattaactaataaggNNNacaaacatgtcaaaaggagaagaacttttt acag |
| | **pRnnnRBS** | ccttattagttaatcgctagcaattgttatccg |
| Insertion of mutation in -10 | **pFnnn-10** | tttatctacaaggtgtggNNNaatgtgtgtaattgtgagcggataacaattg |
| | **pRnnn-10** | ccacaccttgtagataaagtcaacaacttttg |
| Insertion of mutation in -35 | **pFnnn-35** | attttgcaaaagttgttNNNtttatctacaaggtgtggcataatgtgtg |
| | **pRnnn-35** | caacttttgcaaaatgaattgtgagtgctc |
| Insertion of mutation in spacer | **pFnnnSpacer** | ttttgcaaaagttgttgactttatNNNNNNggtgtggcataatgtgtgtaattgtgag |
| | **pRnnnSpacer** | ataaagtcaacaacttttgcaaaatgaattgtgag |
| Primers to amplify pEF1-pHyp in combination with NNN primers | **pFpTRKH2-middle** | acggacacacaactcgatttg |
| | **pRpTRKH2-middle** | caaatcgagttgtgtgtccgt |
